# Phylogenomic analyses support a new infrageneric classification of *Pourthiaea* (Maleae, Rosaceae) using multiple inference methods and extensive taxon sampling

**DOI:** 10.1101/2023.08.13.552906

**Authors:** Guang-Ning Liu, Dai-Kun Ma, Yu Zhang, Richard G.J. Hodel, Si-Yu Xie, Hui Wang, Ze-Tao Jin, Fu-Xing Li, Shui-Hu Jin, Liang Zhao, Chao Xu, Yu Wei, Bin-Bin Liu

## Abstract

In this comprehensive study, we conducted extensive taxon sampling and performed phylogenomic analyses based on plastome and nuclear ribosomal DNA (nrDNA) datasets. We employed multiple inference methods, including concatenated and coalescent-based strategies, to generate an accurate phylogeny of the woody Rosaceae genus *Pourthiaea*. The nrDNA phylogeny of *Pourthiaea* strongly supported three major clades, which were consistent with morphology. However, the plastid tree provided an alternative phylogenetic topology, indicating cytonuclear discordance. Frequent hybridizations between and among the species of *Pourthiaea* could explain the cytonuclear discordance. Considering the evidence from morphology and phylogenomic data, we propose a new infrageneric classification for *Pourthiaea*, consisting of three sections: *P.* sect. *Pourthiaea*, *P.* sect. *Amphidoxae* B.B.Liu, and *P.* sect. *Impressivenae* B.B.Liu.

## INTRODUCTION

The deciduous woody genus *Pourthiaea* Decne. (Maleae, Rosaceae) comprises approximately 11-22 species and is distributed across East Asia to Southeast Asia (Kuan & Yu, 1974; Lu & Spongberg, 2003; Liu & Hong, 2017). It was originally described by the French botanist Joseph Decaisne in 1894 and was named in honor of the Swiss botanist Charles Pourthié. *Pourthiaea* is characterized by deciduous leaves, warty peduncles and pedicels, Kribs’III–I heterogeneous rays in the wood, and clusters of stone cells surrounded by parenchymatous cells in the flesh of pomes, making it easily distinguishable from its allies (Iketani & Ohashi, 1991; Lu & al., 1991; Zhang, 1992). Previously, *Pourthiaea* was considered either a section of *Photinia* Lindl. (Kuan & Yu, 1974; Robertson & al., 1991; Lu & Spongberg, 2003; Campbell & al., 2007; Potter & al., 2007) or a synonym of *Aronia* Medik. (Kalkman, 2004). However, recent phylogenetic and morphological evidence has confirmed its monophyly and generic status (Guo & al., 2011; Liu & al., 2019, 2022).

Despite strong support for the monophyly of *Pourthiaea*, its phylogenetic position remains controversial. Several phylogenetic studies based on whole plastome sequences have suggested a sister relationship between *Pourthiaea* and three species of *Malus* Mill. from different parts of the world, i.e., two Mediterranean species (*M. trilobata* (Labill. ex Poir.) C.K.Schneid. and *M. florentina* C.K.Schneid.) and one species from the New World (*M. ioensis* (Alph.Wood) Britton) (Liu & al., 2019, 2020a, 2020b, 2023). In contrast, various nuclear phylogenies have presented alternative phylogenetic positions of *Pourthiaea*. For instance, some suggested that it is sister to *Docyniopsis* (C.K.Schneid.) Koidz. and *Eriolobus* M.Roem. (GBSSI-1A, Campbell & al., 2007), while others suggested that it forms a clade with *Aronia* and *Aria* (Pers.) Host (nuclear ribosomal ITS (nrITS), Lo & Donoghue, 2012) or with *Cydonia* Mill. (nuclear ribosomal DNA (nrDNA), Liu & al., 2019). Although phylogenomic results of Xiang & al.’s (2017) based on a transcriptomic dataset moderately supported the sister relationship between *Pourthiaea* and a clade including three multi-ovules genera, *Chaenomeles* Lindl., *Cydonia*, and *Pseudocydonia* (C.K.Schneid.) C.K.Schneid., only 16 out of 28 genera in Maleae (Robertson & al., 1991) were sampled in their study. Liu & al. (2022) assembled 797 single-copy nuclear (SCN) genes and provided a robust phylogenomic backbone for the Maleae, in which the sister relationship between *Pourthiaea* and two multi-ovules genera, *Cydonia* and *Pseudocydonia* was strongly supported. However, the tree inferred from 80 plastid coding genes (CDSs) revealed an alternative topology, being sister to the clade II of *Malus*, i.e., the eastern North American and Mediterranean European/West Asia species (Liu & al., 2022, 2023). *Pouthiaea* was concluded to have acted as the maternal parent in a chloroplast capture event associated with the origin of *Malus*. However, this mechanism (i.e., chloroplast capture) requires further investigation in a phylogeographic context.

Yu & Kuan (1963) placed *Pourthiaea* as a section of *Photinia*, designated as *P.* sect. *Pourthiaea*, with two series (ser. *Multiflorae* K.C.Kuan with more than ten flowers and ser. *Pauciflorae* K.C.Kuan with fewer than ten flowers) based on the number of flowers in the inflorescence. However, this infrageneric classification was rejected by recent molecular phylogenetic studies (Guo & al., 2011; Liu & al., 2019). Furthermore, population-level morphological studies (Liu & Hong, 2016a, 2017) have demonstrated that the number of flowers in the inflorescence is highly variable and unsuitable for infrageneric delimitation. Therefore, Guo & al. (2011) proposed the need for an alternative infrageneric classification of *Pourthiaea*.

Species delimitation within *Pourthiaea* has been controversial, largely due to the absence of a comprehensive monograph of the genus across its distribution. Regional floras have documented *Pourthiaea* species, such as the Flora of British India by Hooker (1878), Flora of China by Kuan & Yu (1974) and Lu & Spongberg (2003), and Flore du Cambodge du Laos et du Vietnam by Vidal (1968). However, these taxonomic treatments were based solely on morphological evidence. Liu & Hong (2016a, 2016b, 2017) introduced multivariate statistical analyses and employed the bio-morphological species concept (Hong, 2020) to evaluate character variation between and within populations, and delimitated all taxa into 11 species and one subspecies. However, this method also relied solely on morphological evidence. Phylogenetic relationships among species of *Pourthiaea* demonstrated conflict between plastid and nuclear topologies, i.e., cytonuclear discordance (Guo & al. 2011; Liu & al. 2019). However, the scarcity of molecular data, coupled with a lack of rigorous phylogenetic analyses, has hindered the resolution of taxonomic problems in *Pourthiaea*. Therefore, a robust phylogenomic backbone with comprehensive taxon sampling is essential in resolving taxonomic uncertainties among species.

The taxonomic controversies within *Pourthiaea* may stem from its complicated evolutionary history, possibly including hybridization, polyploidization, apomixis, and incomplete lineage sorting. These evolutionary processes have played essential roles in diversifying other plant genera, such as *Amelanchier* Medik. (Campbell & al., 1985, 1987; Weber & Campbell, 1989; Campbell & Wright, 1996; Dibble & al., 1998; Burgess & al., 2014), *Malus* (Nikiforova & al., 2013; Liu & al., 2022), the expanded *Rhaphiolepis* Lindl. (Liu & al., 2020b; Chen & al., 2022), and *Sorbus* L. (Rich & al., 2010; Robertson & al., 2010; Hamston & al., 2018), as well as the *Amelanchier-Malacomeles* (Decne.) Decne.-*Peraphyllum* Nutt. clade (Liu & al., 2020a).

Genome skimming is a next-generation sequencing (NGS)-based technique that has gained much attention in systematic biology (Coissac & al., 2016). It involves using high-throughput sequencing to generate large amounts of data from low-coverage genome samples, including the complete chloroplast genome and the entire nuclear ribosomal DNA (nrDNA) repeats, typically producing sufficient informative sites for phylogenetic relationships. PCR-free genome skimming (Straub & al., 2012), in particular, has been useful for generating data from herbarium/museum specimens in a cost-effective manner (Baker & Dransfield, 2016; Zeng & al., 2018; Su & al., 2021). This method has been employed to resolve a series of intractable phylogenetic questions in various lineages of Maleae, such as the *Photinia* complex (Liu & al., 2019), the *Amelanchier-Malacomeles-Peraphyllum* clade (Liu & al., 2020a), and *Rhaphiolepis*-*Eriobotrya* Lindl. clade (Liu & al., 2020b). Additionally, the whole plastome has shown great promise in many lineages of angiosperms, such as *Diospyros* L. (Turner & al., 2016), *Fritillaria* L. (Bi & al., 2018), Magnoliaceae (Liu & al., 2020; Wang & al., 2020), *Panax* L. and Araliaceae (Ji & al., 2019; Valcárcel & Wen, 2019), *Theobroma* L. (Kane & al., 2012), and *Vitis* L. and Vitaceae (Zhang & al., 2015, 2016; Wen & al., 2018, 2020; Liu & al., 2021).

In this study, we utilize a genome skimming approach to sequence the entire plastome and the complete nrDNA repeats, with the goal of reconstructing a comprehensive and reliable phylogeny of *Pourthiaea* within the context of Maleae. Furthermore, we intend to propose a novel infrageneric classification based on phylogenomic and morphological evidence.

## MATERIALS AND METHODS

### Taxon sampling, DNA extraction, and sequencing

We sampled 54 individuals in *Pourthiaea* representing all species currently recognized in Lu & Spongberg (2003), Liu & Hong (2016a, 2016b, 2017), and Lou & al. (2022). Our sampling covers the entire range of its distribution (Fig. 1) and includes 51 individuals sequenced for the first time in this study. In order to accurately infer the phylogenetic position of *Pourthiaea* in the framework of the apple tribe Maleae, thirteen genera, represented by 19 species, were selected as outgroups; they are *Amelanchier*, *Crataegus* L., *Cydonia*, *Gillenia* Moench, *Hesperomeles* Lindl., *Kageneckia* Ruiz & Pav., *Lindleya* Kunth, *Malus* Mill., *Phippsiomeles* B.B.Liu & J.Wen, *Photinia*, *Rhaphiolepis*, *Sorbus* L. sensu lato, and *Vauquelinia* Corrêa ex Bonpl. For the widespread species, multiple individuals per species were sampled (Fig. 1 and Appendix 1), including nine individuals of *Pourthiaea amphidoxa* (C.K.Schneid.) Stapf, eight of *P. beauverdiana* (C.K.Schneid.) Hatus., nine of *P. parvifolia* E.Pritz. ex Diels, and six of *P. villosa* (Thunb.) Decne.. Raw sequence data from every individual has been deposited in the NCBI Sequence Read Archive (BioProject PRJNA952323).

**Fig. 1.**
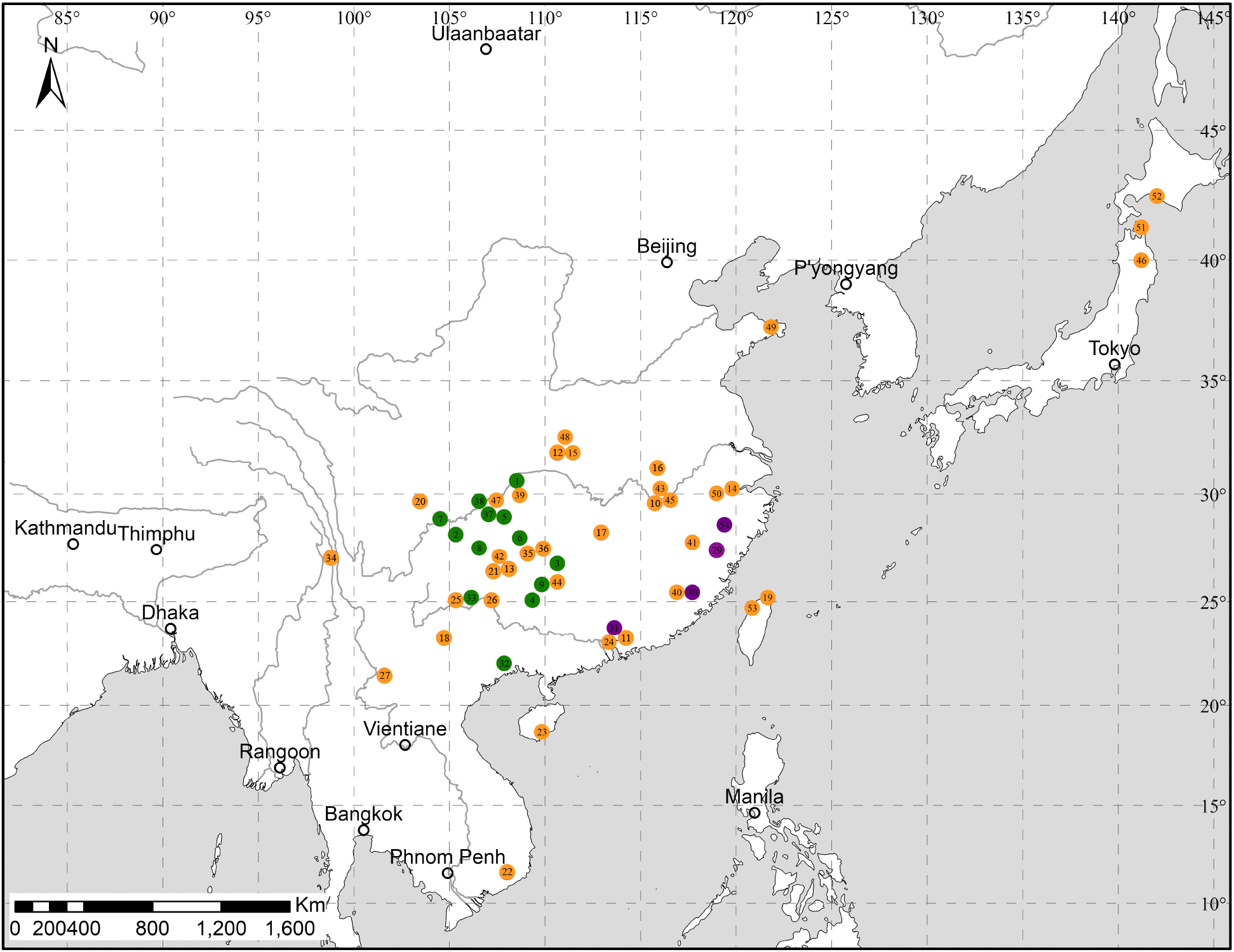
The individuals sampled for this study and the different color corresponds to the clades in the nrDNA tree (Fig. 2A).

Total genomic DNAs were extracted from silica-gel dried leaves and/or herbarium specimens using different protocols in different labs. A modified CTAB (mCTAB) method (Li & al., 2013) was used for the samples indicated by α & β (Appendix 1) in the Institute of Botany, Chinese Academy of Science (IBCAS) in China. The Qiagen DNeasy® plant mini-kit (Qiagen Gmbh, Hilden, Germany) was used for the samples indicated by γ (Appendix 1) following the manufacturer’s protocol in the Laboratories of Analytical Biology (LAB), National Museum of Natural History, Smithsonian Institution in the United States. These samples were sequenced over five years (2017-2022). The libraries of the samples indicated by α (Appendix 1) were prepared in the lab of Institute of Botany, Chinese Academy of Science (IBCAS) in China using the NEXTFLEX® Rapid DNA-Seq Library Prep Kit; the libraries designated by β (Appendix 1) were prepared in the lab of Novogene (Beijing) using NEBNext® Ultra™ II DNA Library Prep Kit for Illumina®; the libraries designated by γ were prepared in the Laboratories of Analytical Biology (LAB), National Museum of Natural History, Smithsonian Institution in the US using NEBNext® Ultra™ II DNA Library Prep Kit for Illumina®. The pooled libraries prepared in China were shipped to Novogene (Beijing) for sequencing, and the pooled libraries in the US were shipped to UC Davis Sequencing Center & CLIA Lab in the USA. Paired-end reads of 2 × 150 bp for all samples were generated on the Illumina HiSeq 2500 Instrument (Novogene Beijing & Novogene America).

### Assembly of the chloroplast genome and nrDNA repeats

A two-step strategy was employed to assemble the plastomes of *Pourthiaea*, and this method allows for obtaining a high-quality chloroplast genome even with low-coverage data. Initially, NOVOPlasty v.4.3.1 (Dierckxsens & al., 2016) was used to assemble the plastomes from high-quality raw data, and the remaining samples were assembled using the **S**uccessive **A**pproach combining **R**eference-based and **D**e novo assembly (hereafter referred to as “**SARD** approach”). This assembly method has been successfully employed in a series of studies, encompassing various angiosperm lineages, such as Amaryllidaceae (Lou & al., 2022), Araliaceae (Liu & Wen, 2019; Liu & al., 2023; Jin & al., 2023), Fabaceae (Su & al., 2019, 2020, 2021), Magnoliaceae (Liu & al., 2020; Wang & al., 2020), Rosaceae (Liu & al., 2019, 2020a, 2020b, 2021, 2022), and Vitaceae (Liu & al., 2021). To facilitate ease of use and citation in future phylogenomic studies, we formalize the SARD approach and provide a detailed step-by-step procedure for assembling the chloroplast genome in the following text.

The program NOVOPlasty is the most accurate and least labor-intensive approach using *de novo* assembly and a seed-and-extend algorithm. However, it requires high-quality raw reads to cover the entire plastome. SARD offers an excellent alternative for obtaining a relatively complete and accurate plastome for lower-quality raw data. However, due to Bowtie 2’s sensitivity to the choice of reference (Langmead & Salzberg, 2012), SARD requires a closely related reference sequence, which in turn requires more time and RAM.

As a first step in running NOVOPlasty v.4.3.1, we removed adapter sequences using cutadapt v.2.7 (Martin, 2011) to trim unwanted adapters with AGATCGGAAGAGC as forward and reverse adapters. Subsequently, we checked the quality of the results with FastQC 0.11.8 (Andrews, 2018) and proceeded to downstream analyses only if all adapter sequences were removed. To increase the likelihood of obtaining a circular chloroplast genome, we selected the ribulose-1,5-bisphosphate carboxylase/oxygenase large subunit (*rbcL*), a plastome-specific and non-mitochondrial sequence of approximately 600 bp, as the seed sequence for the extension. The chloroplast genome of *Pourthiaea villosa* (GenBank accession: MN061989) downloaded from GenBank was used as a reference to correct the direction of inverted repeats. We ran the NOVOPlasty with the other parameters as default. We obtained six (11.5%) circular chloroplast genomes in these 54 samples, including *Pourthiaea amphidoxa* P5 (GenBank accession: MT249055), *P. amphidoxa* P6 (GenBank accession: MT249059), *P. amphidoxa* P7 (GenBank accession: MN061992), *P. blinii* (H.Lév.) Iketani & H.Ohashi P26 (GenBank accession: MN061990), *P. tomentosa* (T.T.Yu & T.C.Ku) B.B.Liu & J.Wen P38 (GenBank accession: MN061995), and *P. villosa* P47 (MN061989). We utilized the SARD approach to assemble the remaining plastomes. Trimmomatic v.0.33 (Bolger & al., 2014) was employed for quality trimming and adapter clipping, followed by quality control evaluation using FastQC 0.11.8 (Andrews, 2018). The SARD approach included two steps: a reference-based method and a *de novo* assembly. For the former, we aligned short reads from six plastomes assembled by NOVOPlasty to the indexed reference using Bowtie 2 (Langmead & Salzberg, 2012) and generated a consensus sequence using Geneious Prime (Kearse & al., 2012). The *de novo* assembly was performed by SPAdes 3.13.1 with careful error correction and a K-mer length of 21, 33, 55, 77 (Nurk & al., 2013). The resulting scaffolds from the *de novo* assembly and contigs from NOVOPlasty were realigned to the draft plastome to correct errors and ambiguities, yielding a high-quality complete plastome. This plastome was subsequently added to the reference database, and a closely related species identified through phylogenetic inference in previous studies was used to assemble subsequent samples, forming the basis of the SARD approach.

The nrDNA repeats are typically present in the plant genome as tandem arrays comprising hundreds or thousands of copies (Poczai & Hyvönen, 2010). We used NOVOPlasty v. 3.7.2 to assemble the entire nrDNA repeats. Adapter sequences were also trimmed using cutadapt v.2.7 (Martin, 2011) and quality controlled by FastQC 0.11.8 (Andrews, 2018). We obtained three complete sequences of nrDNA repeats, *Pourthiaea sorbifolia* (W.B.Liao & W.Guo) B.B.Liu & D.Y.Hong P36, *P. villosa* P47, and *P. villosa* P50. Using the SARD approach, these sequences were then used as a reference to assemble the remaining samples. Due to the inherent variability in the structure of the external transcribed spacers (ETS), assembling the complete ETS region was not feasible. However, despite this challenge, we retained the conserved portion of the ETS among species to ensure its inclusion in the subsequent phylogenomic analyses. By retaining this conserved segment, we aimed to capture valuable evolutionary information that can contribute to our understanding of the relationships and evolutionary history within the studied taxa.

### Plastome and nrDNA annotation

The assembled plastid genomes were annotated using Geneious Prime (Kearse & al., 2012), with the closest related plastome downloaded from GenBank (*Pourthiaea villosa*: accession no. MN061989) as a reference. The annotation of the coding sequences was manually checked by translating them into proteins to verify the start and stop codons in Geneious Prime. To determin the boundaries of the large-single copy (LSC), small-single copy (SSC), and inverted-repeats (IRs), the Find Repeats tool in Geneious Prime was utilized, considering the presence of two reverse complementary repeats observed in the plastomes of species in the Rosaceae. The custom annotations were then converted to the required format for NCBI submission, including FASTA and five-column feature tables, using GB2sequin (Lehwark & Greiner, 2019). Lastly, the gene map of *Pourthiaea* was generated using OrganellarGenomeDRAW (OGDRAW) version 1.3.1 (Greiner & al., 2019).

### Data matrix generation and sequence cleaning

The entire nrDNA sequence includes six independent regions, ETS, 18S, Internal Transcribed Spacer 1 (ITS1), 5.8S, Internal Transcribed Spacer 2 (ITS2), and 26S. All these six nuclear regions were concatenated as “nrDNA dataset”. Due to the missing data resulting from sequence divergence and potential poor alignment, we generated four plastid data matrixes with various species sampling strategies and distinct locus coverage for accurate phylogenetic inference: 1) whole plastome data matrix with all 54 *Pourthiaea* samples and only one outgroup, *Cydonia oblonga* Mill. (referring to as “WP55 dataset” hereafter); 2) whole plastome data matrix with all 73 individuals sampled in the framework of Maleae (referring to as “WP73 dataset” hereafter); 3) 79 plastid concatenated CDSs with all 54 *Pourthiaea* samples and only one outgroup, *Cydonia oblonga* Mill. (referring to as “CDS55 dataset” hereafter); 4) 79 plastid concatenated CDSs with all 73 individuals sampled in the framework of Maleae (referring to as “CDS73 dataset” hereafter). All these five data sets are available from the Dryad Digital Repository https://doi.org/10.5061/dryad.np5hqc001

The six regions of the nrDNA repeats and 79 plastid CDSs were aligned separately by MAFFT v. 7.520 (Nakamura & al., 2018) with options “—localpair –maxiterate 1000”. The aligned nuclear nrDNA and plastid CDSs will be concatenated by AMAS (Borowiec, 2016) and directly used for the following phylogenetic inference. As the sequences of the two IR regions of the whole plastome in each assession of the Maleae were completely or nearly identical, only one copy of the inverted repeat (IR) region was included for the whole plastome phylogenetic analyses. To reduce systematic errors produced by poor alignment, we used trimAL v1.2 (Capella-Gutiérrez & al., 2009) to trim the alignment of the plastome. We, furthermore, employed Spruceup (Borowiec, 2019) to discover, visualize, and remove outlier sequences with window size 50 and overlap 25.

### Integrating multiple methods for accurate phylogenetic inference

The best-fit partitioning schemes and nucleotide substitution models for the unpartitioned whole plastome datasets, partitioned plastid CDSs datasets, and partitioned nrDNA dataset were estimated using PartitionFinder2 (Lanfear & al., 2016; Stamatakis, 2006). Under the corrected Akaike information criterion (AICc) and linked branch lengths, the PartitionFinder2 was performed using the greedy setting (Lanfear & al., 2012) for plastid WP55 and WP73 datasets, rcluster (Lanfear & al., 2014) algorithm option for plastid CDS55, CDS73, and nuclear nrDNA datasets with prior defined data blocks by codon positions of each protein-coding genes and all models. The partitioning schemes and evolutionary model for each subset were used for the downstream Maximum Likelihood (ML) and Bayesian Inference (BI) analyses. The ML tree was inferred by IQ-TREE2 v.2.1.3 (Nguyen & al., 2015; Minh & al., 2020) with 1000 SH-aLRT and the untrafast bootstrap replicates using UFBoot2 (Hoang & al., 2017), with the collapse near-zero branches option. We also used an alternative ML tree inference method, RAxML v. 8.2.12 (Stamatakis, 2014) with GTRGAMMA model for each partition and clade support assessed with 200 rapid bootstrap (BS) replicates. The BI was performed with MrBayes 3.2.7 (Ronquist & al., 2012). The Markov chain Monte Carlo (MCMC) analyses were run for 50,000,000 generations. Trees were sampled at every 1,000 generations with the first 25% discarded as burn-in. The remaining trees were used to build a 50% majority-rule consensus tree. The stationarity was regarded to be reached when the average standard deviation of split frequencies remained below 0.01. Each ML gene tree of 79 plastid coding genes was estimated by RAxML 8.2.12 (Stamatakis, 2014) with a GTRGAMMA mode and the option “-f a” and 200 BS replicates to assess clade support. The resulting ML trees were used to estimate a coalescent-based species tree with ASTRAL-III (Zhang & al., 2018) with local posterior probabilities (LPP; Sayyari & Mirarab, 2016) to assess clade support. Given the effect of the low support branches, we rooted each plastid CDS gene tree and collapsed ≤ 10% support branches by *phyx* (Brown & al., 2017). All the ML (ten), BI (five), and ASTRAL species trees (two) were visualized using Geneious Prime (Kearse & al., 2012), and all 17 trees have been deposited in the Dryad Digital Repository https://doi.org/10.5061/dryad.np5hqc001

## RESULTS

### Phylogenetic analyses of nrDNA sequences

The nrDNA repeats included six regions: partial ETS, 18S, ITS1, 5.8S, ITS2, and 26S. The aligned matrix was 6332 bp in length, with poorly aligned regions trimmed. To ensure accurate phylogenetic inference, three trees were generated, two ML trees (RAxML and IQ-TREE2) and one BI tree (MrBayes). All three phylogenies provided strong support for three distinct clades (Fig. 2 and suppl. Figs. S2-S4): clade A (BS/SH-aLRT/Ultrafast BS/BI: 100/100/100/1), clade B (93/98/95/1), and clade C (82/99/100/1). Clade A, positioned as the basalmost clade, encompassed three species: *Pourthiaea hirsuta*, *P. impressivena* (Hayata) Iketani & H.Ohashi, and *P. zhejiangensis* (P.L.Chiu) Iketani & H.Ohashi; all three species were supported to be monophyletic. Clade B also included three currently recognized species: *P. amphidoxa*, *P. pilosicalyx*, and *P. tomentosa*, all of which formed monophyletic groups. The remaining species were placed within clade C. This clade comprised the widespread species *P. arguta* (Lindl.) Decne. and *P. beauverdiana*, as well as three narrowly distributed species, *P. blinii*, *P. dabeishanensis*, *P. sorbifolia*, and *P. tsaii* (Rehder) Iketani & H.Ohashi, all of which formed a subclade. The other subclade includes two widely distributed species, *P. villosa* and *P. parvifolia*, along with three endemic species: *P. magnoliifolia* (Zhejiang, China), *P. lucida* Decne. (Taiwan, China), *P. pustulata* (Guangdong & Hainan, China).

**Fig. 2.**
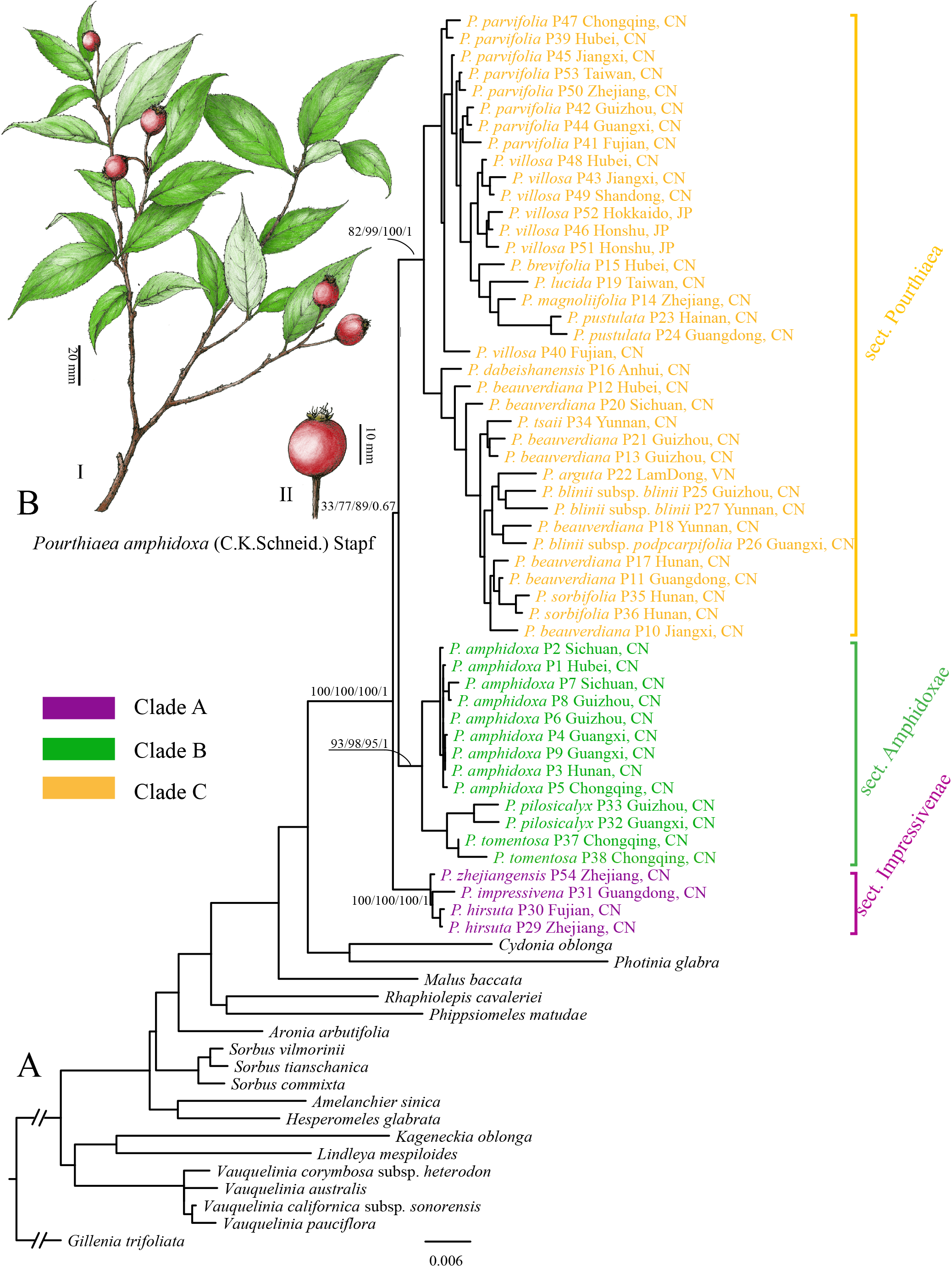
**A**: Maximum likelihood phylogeny of *Pourthiaea* in the framework of Maleae inferred from RAxML analysis of the nuclear ribosomal DNA (nrDNA) data matrix. The numbers near the five focused nodes are the Bootstrap Support (BS) inferred from RAxML, the SH-aLRT support and Ultrafast Bootstrap support inferred from IQ-TREE2, and BI posterior probabilities inferred by MrBayes (from left to right), respectively. **B**: an illustration of *Pourthiaea amphidoxa*.

### Phylogenetic relationships based on chloroplast genome

The 54 chloroplast genomes of *Pourthiaea*, which encompassed all currently recognized species, exhibited a typical quadripartite structure similar to other published plastomes in Maleae with lengths from 160,142 to 160,496 bp (suppl. Fig. S1). They consisted of two inverted repeats (IRs) that were complementary to each other, with a small single copy (SSC) and a large single copy (LSC) separating them (suppl. Fig. S1). In total, the plastomes encoded 132 genes, including 85 coding genes, 37 tRNA genes, and eight rRNA genes (suppl. Fig. S1).

The aligned plastome matrix used for ML and BI analysis had a length of 132,929 bp, with regions of pool alignment trimmed. We generated a total of six trees based on the whole plastome datasets (WP55 and WP73) using three phylogenetic inference methods (ML and BI). Additionally, we generated eight trees based on the CDSs datasets (CDS55 and CDS73) using four phylogenetic inference methods (ML, BI, and species tree). These analyses resulted in a total of 14 trees (Fig. 3B and suppl. Figs. S5-S18), all of which recovered two well-supported clades, designated as clade I and clade II. Clade I included a subset of individuals from *P. amphidoxa* (3/9) and *P. pilosicalyx* (T.T.Yu) Iketani & H.Ohashi (1/2), and all 14 trees supported the sister relationship between these two species. Notably, most species with multiple accessions (e.g. *P. amphidoxa*, *P. beauverdiana*, *P. pilosicalyx*, *P. hirsuta* (Hand.-Mazz.) Iketani & H.Ohashi, and *P. villosa*) exhibited polyphyly in the plastome tree, with the exception of the paraphyletic *P. blinii*. Only one species, *P. tomentosa*, was consistently recovered as monophyly.

**Fig. 3.**
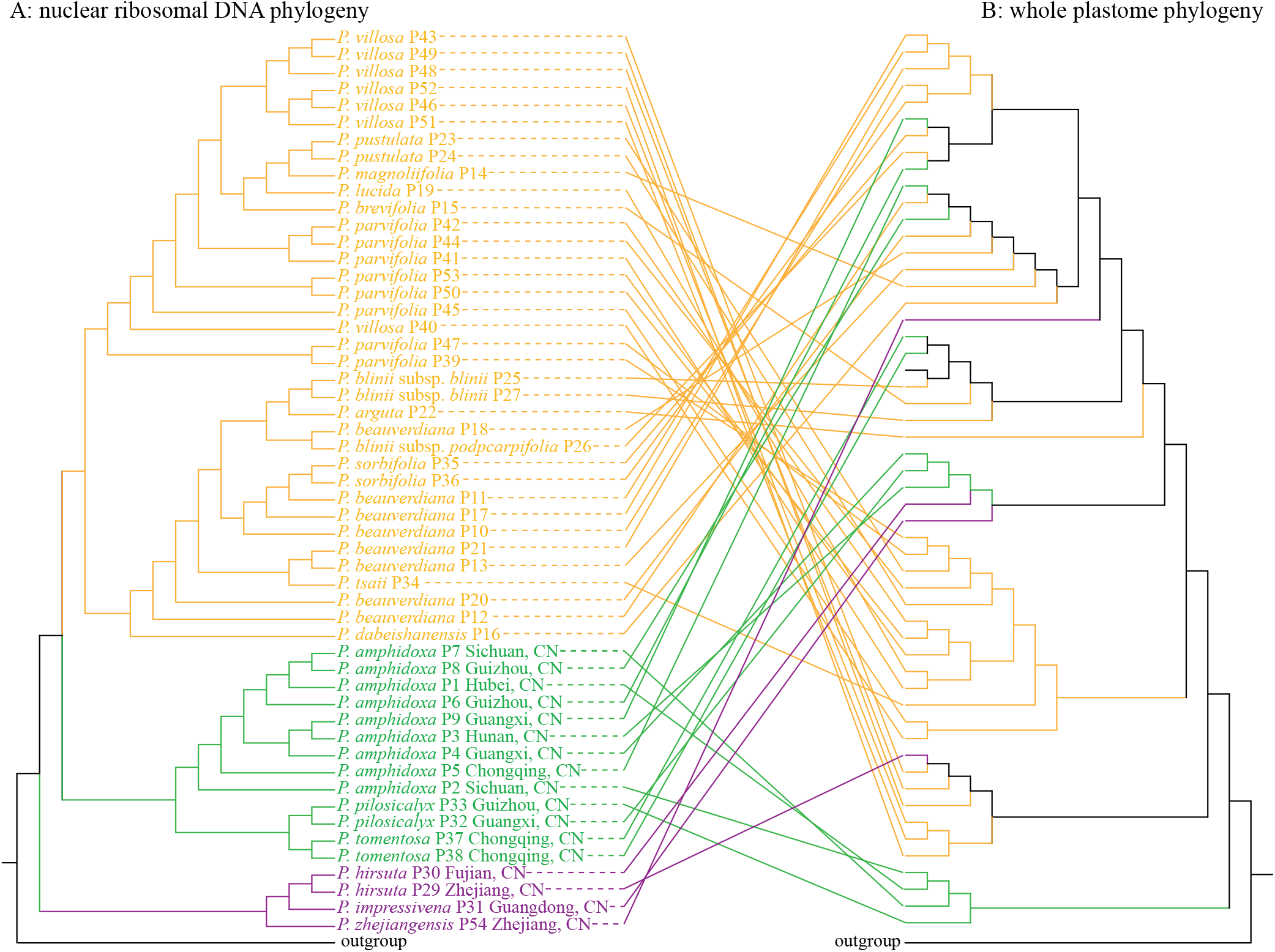
COMPARISONS of the nuclear ribosomal DNA (nrDNA) repeats (left: A) and whole plastome (rights: B) phylogenies, and the color of each one corresponds to the clades in the nrDNA tree (Fig. 2A).

## DISCUSSION

We have generated a substantial dataset comprising 54 individuals in the *Pourthiaea* ingroup and 19 outgroup species in the Maleae. For this study, we successfully assembled the whole plastome and nrDNA repeats. We employed various inference methods to ensure accurate phylogeny, including ML, BI, and a coalescent-based species tree. Each method has its own strengths and assumptions, encompassing a wide range of approaches. These methods rely on specific assumptions regarding evolutionary processes, such as substitution models, rates of evolution, and tree topologies. By utilizing multiple methods, we were able to validate and reinforce the results obtained from each approach. Our findings consistently indicated that these diverse methods yield similar phylogenetic trees, revealing three major clades in our nrDNA tree and two clades in the plastid tree. This outcome substantially bolstered our confidence in proposing a three-section infrageneric taxonomy system.

### Cytonuclear discordance and its potential mechanisms

We observed incongruence between the plastid and nrDNA topologies in *Pourthiaea* (Fig. 3). Various mechanisms could account for this incongruence, including reticulation processes such as allopolyploidy, hybridization, chloroplast capture, and introgression, as well as incomplete lineage sorting or paralogs of nrDNA (Rieseberg & Soltis, 1991). However, limited cytological studies have reported no instances of allopolyploidy in *Pourthiaea*. Furthermore, although polyploidy is common in larger Maleae genera like *Amelanchier* and *Crataegus* L. (Campbell & al., 1991), it has never been documented in this particular genus. Additionally, the scarcity of degenerate bases in nrDNA sequences, representing polymorphisms at specific loci, suggests that incomplete concerted evolution resulting from polyploidy is likely not an explanation for the observed topology. Moreover, our well-supported nrDNA tree, which coincides with morphological characters of the monophyletic species, excludes the possibility of biparental nuclear genes contributing to the high-resolution topology. This further eliminates allopolyploidy as a potential cause. While incomplete lineage sorting can also lead to conflict between nuclear and plastid topologies, distinguishing between hybridization and lineage sorting poses challenges (Joly & al., 2009). It is worth noting that although the concerted evolution of nrDNA arrays within individuals may result in homogeneity (Bailey & al., 2003), intra-individual polymorphism has been observed in certain lineages with hybrid origins (Xu & al., 2017). Nevertheless, as nrDNA paralogy did not impede the accurate reconstruction of the phylogenetic relationships among *Pourthiaea* species, it is unlikely to explain the incongruence between the nuclear and plastid topologies.

Hybridization is the most likely explanation for the cytonuclear discordance observed in *Pourthiaea*. In the Maleae, hybridization has been widespread both within and between genera (Robertson & al., 1991). Some genera, such as *Micromeles* and *Pseudocydonia* (Lo & Donoghue, 2012), as well as *Phippsiomeles* (Liu & al., 2019), have been proposed to have hybrid origins. Hybridization events have also been identified within the expanded genus *Rhaphiolepis* (Liu & al., 2020b). Additionally, chloroplast capture has played a significant role in the evolution of *Malacomeles* and *Peraphyllum* (Liu & al., 2020a). Therefore, extensive hybridization may have contributed to the diversification of *Pourthiaea* (Fig. 3). For instance, *Pourthiaea hirsuta* P29, which occurs in Zhejiang (China), may have captured the chloroplast genome of *P. villosa* (Figs. 1-2), with the most recent common ancestor (MRCA) of *P. hirsuta* serving as the paternal parent and the MRCA of *P. villosa* as the maternal parent. According to the pollen competition scenario (Rieseberg, 1995), hybridization mostly occurred in the MRCA population of *P. hirsuta*, where individuals of this species provided pollen for the other species (*P. villosa*) in the minority (Rieseberg, 1995; Acosta & Premoli, 2010; Liu & Hong, 2017). The wide distribution of pome fruits of *Pourthiaea*, ranging from Japan to China (Liu & Hong, 2016a), makes them a common food souce for small animals, including birds with long-distance migratory patterns. This could have led to the dispersal of *P. villosa* seeds and contributed to their presence in the distribution of other species.

### Phylogenetic implications based on the strongly supported backbone

The monophyly of three major clades in the nrDNA tree (Fig. 2 and suppl. Figs. S2-S4) lacks support in our plastid tree (Fig. 3, and suppl. Figs. S5-S18), revealing significant cytonuclear discordance. Frequent hybridization events have had a substantial impact on the monophyly of some species, which was represented in our nrDNA tree (Fig. 2). We will present additional morphological evidence for these three major clades in our nrDNA topology.

### The Pourthiaea impressivena clade (Clade A)

The *Pourthiaea impressivena* clade is comprised of three species recognized in previous studies, *P. hirsuta*, *P. impressivena*, and *P. zhejiangensis* (Fig. 2). *P. impressivena*, characterized by more than ten flowers in the inflorescences and previously considered distantly related to *P. hirsuta* and *P. zhejiangensis* (Kuan & Yu, 1974; Lu & Spongberg, 2003). However, our nrDNA phylogeny robustly establishes a close relationship among these three species; *P. hirsuta* and *P. impressivena* form a clade, which is then sister to *P. zhejiangensis*. The close relationship among these three species is supported by a shared character, the ferruginous villous pubescence on the leaf blade, petiole, inflorescences, hypanthium, and branchlet (Fig. 4E). In the plastid tree, these three species are distributed across the distinct clades (Fig. 3B), and two individuals of *P. hirsuta* were found to be non-monophyletic.

**Fig. 4.**
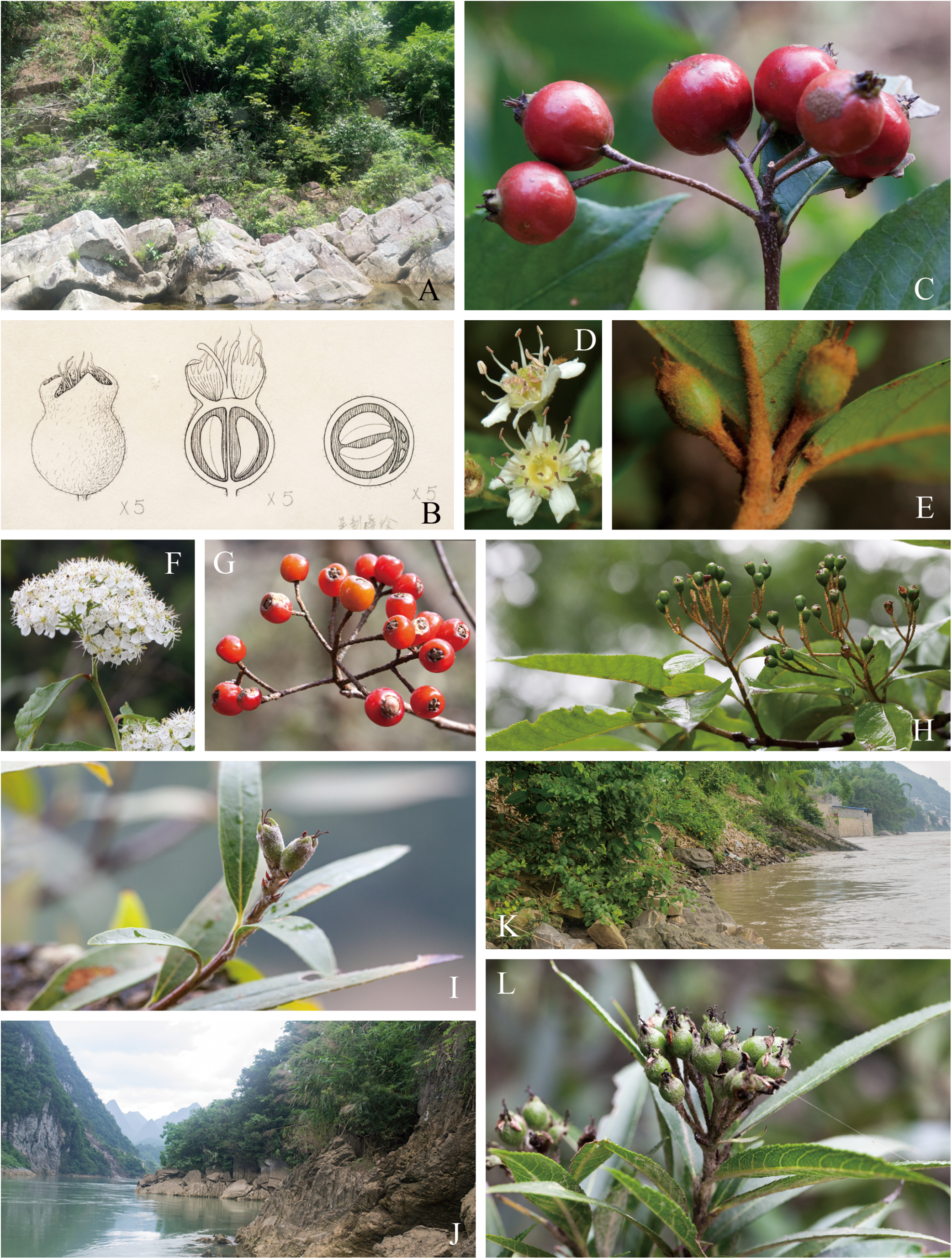
Representative species in *Pourthiaea*. ***P. pilosicalyx***: A, the habitat; B, the urceolate fruits and two carpels in one ovary; ***P. amphidoxa***: C, large fruits (larger than 10 mm in diameter), D, five carpels in one ovary; ***P. hirsuta***: E, the ferruginous villous leaf blade, petiole, inflorescences, hypanthium, and branchlet; ***P. beauverdiana***: F, inflorescences; G-H, mature and young fruits; ***P. blinii***: I, young fruits, J, habitat; ***P. tsaii***: K, habitat, L, young fruits. Photos credit: A (Guangxi, China), C (Hunan, China), F (Guizhou, China), G (Hunan, China), H (Zhejiang, China), I (Yunnan, China), J (Guangxi, China), K (Yunnan, China), and L (Yunnan, China) to Bin-Bin Liu; B (PE [barcode 00934236]); D (Sichuan, China) to Wen-Bin Ju; E (Zhejiang, China) to Cheng-Chun Pan.

The incongruence between the plastid and nuclear topologies in this clade can be attributed to the potential hybridization events that occurred during its evolution. Considering the cytonuclear conflicts in *Pourthiaea zhejiangensis*, our data support the hypothesis that the ancestral lineages of *P. zhejiangensis* likely served as the maternal parent. In contrast, the paternal parent may be a member of clade I (Fig. 3B). *Pourthiaea zhejiangensis* exhibits similarities to *P. hirsuta*, such as ferruginous branchlets, leaf blade, peduncle, hypanthium, and corymb inflorescences (Fig. 4E), while also sharing traits with *P. beauverdiana*, including conical winter buds and thick leaf blades. Furthermore, *P. zhejiangensis* is endemic to the southern regions of Zhejiang, China, suggesting that this hybrid species may present an ecotype adapted to the low-altitude mountains in the subtropical regions of China. Distinguishing characteristics of *P. zhejiangensis* include slender pedicels (in contrast to shorter pedicels), fewer flowers (1 or 2, rarely 3-6, as opposed to 3-8), and an acute to shortly acuminate leaf apex (versus an acuminate to caudate leaf apex) (Qiu, 1980; Lu & Spongberg, 2003). Based on these distinctions, we hereby recognize *P. zhejiangensis* as a separate species within this clade.

*Pourthiaea impressivena* is the sister species to *P. hirsuta* in both the plastid and the nuclear DNA phylogenies (Figs. 2, 3). *P. impressivena* can be distinguished from *P. hirsuta* by its deeply impressed leaf veins. Additionally, these two species exhibit allopatric distributions, with *P. hirsuta* inhabiting Central to East China, while *P. impressivena* is found in South China.

### The Pourthiaea amphidoxa clade (Clade B)

The *Pourthiaea amphidoxa* clade stands out from the other two clades, characterized by its two or five carpels (rarely four; Fig. 4D) and short petioles. This clade primarily occupies regions from central to South China (Fig. 1). *P. pilosicalyx*, with two carpels in each ovary (versus five carpels, rarely four, Fig. 4B,D), caducous sepals (versus persistent), and urceolate fruits (versus ovoid or globose, Fig. 4B,C), can be readily distinguished from the other two species. Moreover, *P. pilosicalyx* is typically found in riparian habitats (Fig. 4A) and is distributed from Southwest Guizhou (China) to north Vietnam. In contrast, the other two species mainly inhabit forest edges and/or slopes in central China (Liu & Hong, 2017). Two samples were sequenced for this study, one from Guizhou, China, and the other from Guangxi, China. The latter formed a clade with *P. amphidoxa* from Guangxi, suggesting the possibility of hybridization between the ancestral lineages of these two species, with the ancestor of *P. pilosicalyx* capturing the chloroplast genome of *P. amphidoxa*.

Cytonuclear discordance was detected based on the phylogenetic position of *Pourthiaea tomentosa* (Figs. 2, 3 and suppl. Figs. S2-S18). Our nrDNA phylogeny supported a sister relationship between *Pourthiaea tomentosa* and *P. amphidoxa* (Fig. 2), while these two individuals were embedded within three samples of *P. blinii* in the plastid topology (Fig. 3B). Morphological evidence also supported the close relationship between these three species. *P. tomentosa* shared the characteristic of having four or five carpels in one ovary with *P. amphidoxa*, as well as tomentose branchlets, petioles, adaxial leaf blade, peduncles, pedicels, and sepals with *P. blinii*. In our previous study, this conflict was similarly identified and explained as a result of hybrid origin (Liu & al., 2019). The cytonuclear discordance can be attributed to chloroplast capture through multiple instances of hybridization and recurrent backcrossing events. Building on the hypothetical scenario of chloroplast capture (following Acosta & Premoli, 2010; Liu & al., 2020a), we conclude that the ancestral lineages of *P. tomentosa*, serving as the paternal parent, captured the chloroplast genome of the MRCA of *P. blinii*, acting as the maternal parent. Furthermore, *P. tomentosa* is endemic to Chongqing (Jinyunshan and Jinfoshan), China, which may have served as a museum/sanctuary for this species, similar to the case of *Metasequoia glyptostroboides* Hu & W.C.Cheng.

*Pourthiaea amphidoxa* can be easily distinguished from *P. tomentosa* by its large fruits (Figs. 2B-II, 4C) measuring more than 10 mm in size, and its villous or glabrous inflorescences (versus tomentose inflorescences). This distinction has been noted by Lu & Spongberg (2003). Due to the weak reproductive isolation between *P. amphidoxa* and other species within *Pourthiaea* genus, it is possible that *P. amphidoxa* has undergone hybridization events with *P. beauverdiana*, *P. blinii*, *P. hirsuta*, and *P. sorbifolia* in different locations, potentially acting as a maternal parent.

### The *Pourthiaea villosa* clade (Clade C)

There are significant taxonomic controversies surrounding this clade. Previous regional floras recognized ten species and eight varieties (Ohwi, 1965; Kuan & Yu, 1974; Lu & Spongberg, 2003). However, a recent taxonomic revision of *Pourthiaea* has recognized only two species and one subspecies (Liu & Hong, 2016a, 2017). The nrDNA tree establishes the monophyly of the *Pourthiaea villosa* clade, characterized by shrubs, occasionally small trees, corymbose or umbellate inflorescences, and shorter petioles (Fig. 5). However, these samples are distributed across four clades in the plastid phylogeny (Fig. 3).

**Fig. 5.**
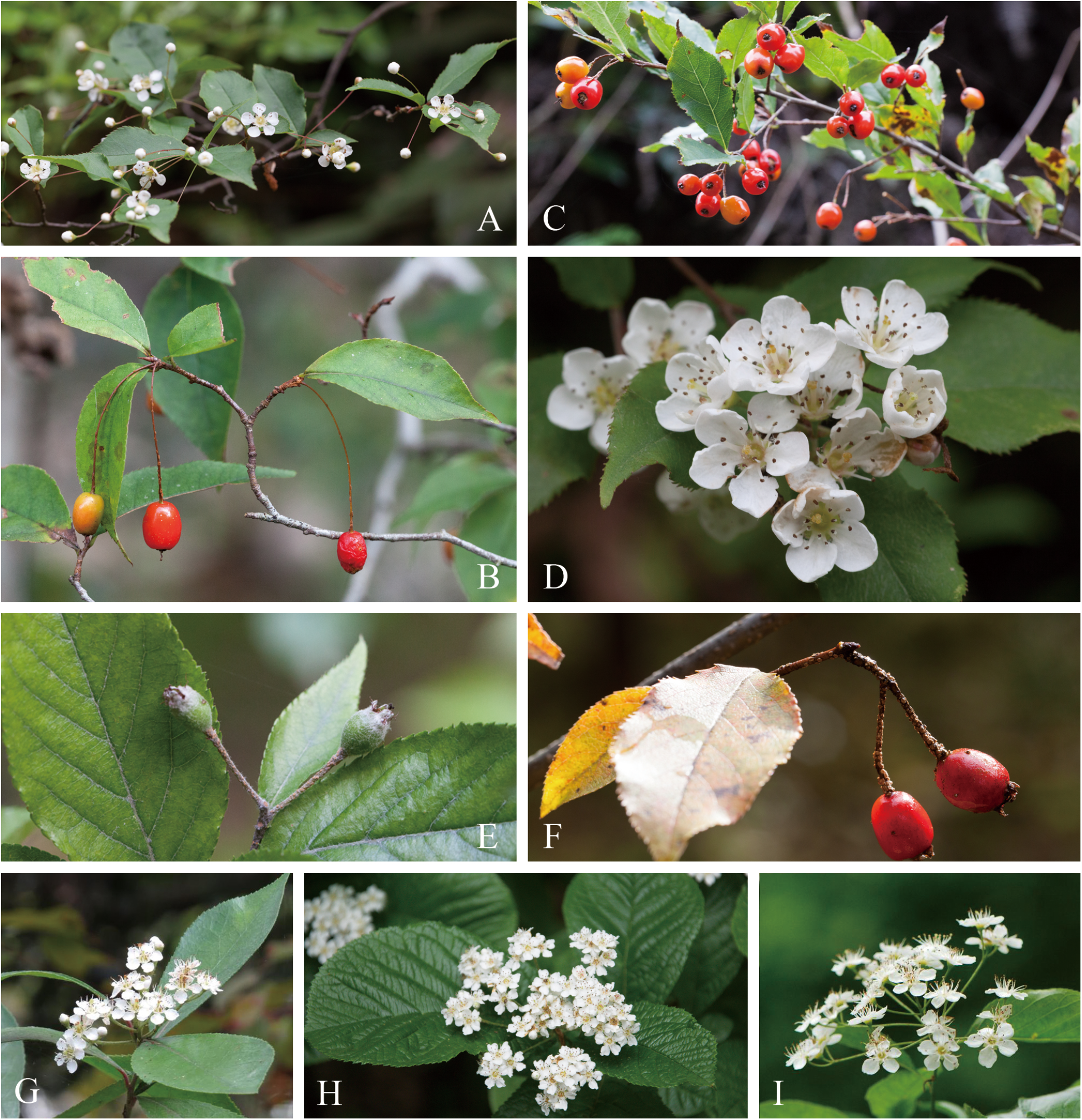
Representative species in *Pourthiaea**. P. komarovii***: A, inflorescences, B, fruits; ***P. parvifolia***: C, fruits, D, inflorescences; ***P. villosa***: E-F, fruits; ***P. pustulata***: G, fruits; ***P. magnoliifolia***: H; ***P. brevifolia***: I. Photo credit: A (Guizhou, China), B (Zhejiang, China), C (Hunan, China), D (Guizhou, China), E (Jiangxi, China), F (Hubei, China), and H (Zhejiang, China) to Bin-Bin Liu; G (Hongkong, China) to Jin-Long Zhang; I (Jiangsu, China) to Rui Tang.

*Pourthiaea villosa* and *P. parvifolia* are represented as distinct clades in the plastid and nrDNA topologies, respectively. *P. parvifolia* (Fig. 5A-D) could be easily distinguished from *P. villosa* (Fig. 5E,F) by its glabrous or slightly villous (versus villous) branchlets, leaf blades, petioles, inflorescences, and hypanthium. The inflorescences of *P. parvifolia* form a corymb, occasionally a compound corymb (versus compound corymb or umbel, rarely corymb). Furthermore, the length of petioles is typically fewer than 4 mm (versus more than 4 mm) in *P. parvifolia*. Additionally, *P. villosa* is primarily distributed in high latitudes and at high altitudes in low latitudes, while *P. parvifolia* inhabits the low latitudes of subtropical China. These two species exhibit distinct allopatric distributions. In Taiwan, China, one population of *P. parvifolia* is characterized by decumbent shrubs with a highly branching habit, and it thrives in sunny gravelly ridges of limestone in Mt. Chingshui, from 600 to 2,100 m in elevation.

The phylogenetic positions of *Pourthiaea brevifolia* (Fig. 5I), *P. lucida*, *P. magnoliifolia* (Fig. 5H), and *P. pustulata* (Fig. 5G) exhibit incongruence between the chloroplast genome and nrDNA phylogenies. This incongruence may be explained by the potential hybridization events that occurred between species. For instance, considering *P. magnoliifolia* as an example, *P. beauverdiana*, which is likely the maternal parent, had a widespread distribution in Zhejiang, China, overlapping with the ancestral range of *P. magnoliifolia*. This sympatric distribution provided an opportunity for hybridization between the two species. *Pourthiaea magnoliifolia* may have originated from hybridization, with *P. beauverdiana* serving as the maternal parent and the MRCA of *P. magnoliifolia* acting as the paternal parent. Additionally, *P. magnoliifolia* shares several morphological characteristics with *P. beauverdiana*, such as a large compound corymb and small tree size. It is possible that the other three species, *P. brevifolia*, *P. lucida*, and *P. pustulata*, also originated from similar hybrid processes.

Our nrDNA phylogeny strongly supports the monophyly of the *Pourthiaea arguta* clade (Fig. 2); however, the samples in the plastid phylogeny (Fig. 3B) form two separate clades. All members within this clade are characterized by their tree-like growth habit, compound corymbose inflorescences, and longer pedicels, setting them apart from other species within the genus *Pourthiaea*. Based on the available morphological evidence, the number of recognized species within this clade varies from four (Liu & Hong, 2017) to 10 (Kuan & Yu, 1974; Lu & Spongberg, 2003; Vidal, 1968).

Three specimens of *Pourthiaea blinii* (P25, P27, and P28) exhibited distinct phylogenetic positions in the plastid topology (Fig. 3B), whereas they are nested within *P. beauverdiana* in the nrDNA tree (Fig. 2). The cytonuclear incongruence observed in these three samples of *P. blinii* could potentially be attributed to hybridization events. Additionally, our field observations revealed that individuals within a population were nearly identical, with only slight character variations (Liu & al., 2016b). According to our field observations in southwest China, *P. blinii* predominantly inhabits riverbank and may experience flooding during the rainy season (Fig. 4J). Subsequently, these populations have adapted to the challenging habitat in the hot valley of the Hongshui River in southwest China, exhibiting adaptive traits such as dwarf shrubs, clumps of stems, and small, thick leaf blades without conspicuous petioles (Fig. 4I: Lo & Donoghue, 2012; Liu & Hong, 2016b). One sample of *P. blinii* from Guangxi, China did not form a clade with other samples of *P. blinii* from Guizhou and Yunnan in both the plastid and nrDNA trees. Furthermore, the Guangxi population was characterized by glabrescent inflorescences, branchlets, and leaves (Fig. 4I), which easily distinguished it from the populations in Guizhou and Yunnan, where inflorescences, branchlets, and leaves were tomentose.

While the sample of *Pourthiaea dabeishanensis* was nested within the clade of *P. beauverdiana* in the plastid phylogeny (Fig. 3B), our nrDNA tree places it as the basalmost position within one subclade of clade C with moderate support (Fig. 2). *P. dabeishanensis* has been previously considered as a synonym of *Photinia schneideriana* (Lu & Spongberg, 2003) due to its glabrescent inflorescences. However, we still need to assemble more nuclear genes to perform robust phylogenomic analyses to confirm the unique phylogenetic position of *P. dabeishanensis*.

*Pourthiaea beauverdiana*, a widely distributed species in subtropical China, has consistently been recognized as a distinct species at the species level (Kuan & Yu, 1974; Hiroyoshi, 1993; Lu & Spongberg, 2003). It can be distinguished from other species by a set of diagnostic characteristics, such as tree-like growth habit, compound corymb inflorescences, and long petioles (Fig. 4F,G,H). However, our nrDNA phylogeny strongly supports the polyphyly of *P. beauverdiana*. Several factors may contribute to this outcome. Frequent hybridization events between species, particularly within relatively young lineages, have led to polymorphisms in nuclear genes. Furthermore, incomplete concerted evolution of nrDNA could also contribute to the observed polyphyly. Additionally, allopolyploidization and subsequent apomixis may further contribute to the polyphyly of *P. beauverdiana*. Pending further study, we provisionally maintain the species status of *P. beauverdiana*, recognizing the need for additional investigations.

### A new infrageneric classification of *Pourthiaea*

Yu & Kuan (1963) initially classified *Pourthiaea* as a section of *Photinia*, *Photinia* sect. *Pourthiaea*. Within this section, they further subdivided it into two series based on the number of flowers in the inflorescence: ser. *Multiflorae* (more than 10 flowers) and ser. *Pauciflorae* (fewer than 10 flowers). However, recent molecular phylogenetic studies utilizing two chloroplast markers plus nrITS (Guo & al., 2011) and the whole plastome and entire nrDNA repeats (Liu & al., 2019) have refuted this morphology-based infrageneric classification. Additionally, morphological studies conducted at the population level (Liu & al., 2016a; Liu & Hong, 2017) have demonstrated significant variability in the number of flowers within the inflorescence. For instance, in the case of *Pourthiaea villosa*, the inflorescence can vary from a corymb or compound corymb to a sub-umbel or compound sub-umbel, and the number of flowers can range from one to 23 (Fig. 5: Liu & al., 2016a).

The taxonomic issues in the genus *Pourthiaea* have broader implications for conservation and biodiversity studies. Accurate species identification is crucial for understanding the distribution, ecology, and evolutionary history of organisms. Furthermore, conservation efforts rely on effectively distinguishing and prioritizing endangered or threatened species. Therefore, resolving the taxonomic uncertainties in *Pourthiaea* is not only of scientific interest but also has practical implications for managing and preserving biodiversity. Here we propose a new infrageneric classification based on our nrDNA results, in which three clades in *Pourthiaea* were strongly supported, i.e., the *P. amphidoxa* clade, the *P. impressivena* clade, and the *P. villosa* clade.

***Pourthiaea*** Decne. Nouv. Arch. Mus. Par. Ser. I, x. 146. 1874. Lectotype, designated by Iketani & Ohashi (1991): *Pourthiaea villosa* (Thunb.) Decne., Nouv. Arch. Mus. Par. Ser. I, x. 147. 1874.

Diagnosis: Deciduous leaves, warty peduncles and pedicels, Kribs’III–I heterogeneous rays in the wood, and clusters of stone cells surrounded by parenchymatous cells in the flesh of pomes.

Distribution: Bangladesh, China, India, Japan, Laos, Myanmar, North Korea, South Korea, Thailand, and Vietnam.

I. ***Pourthiaea*** sect. ***Pourthiaea*** Type: Pourthiaea villosa (Thunb.) Decne.

Diagnosis: This section was characterized by the whitish villous or tomentose leaf blade, petiole, inflorescences, hypanthium, and branchlet.

Distribution: China, India, Japan, Laos, Myanmar, South Korea, and Vietnam. Including ca. 12 species, *Pourthiaea arguta*, *P. beauverdiana*, *P. blinii* subsp. *blinii*, *P. blinii* subsp. *podocarpifolia*, *P. brevifolia*, *P. dabeishanensis*, *P. lucida*, *P. magnoliifolia*, *P. parvifolia*, *P. pustulata*, *P. sorbifolia*, *P. tsaii*, and *P. villosa*.

Five new combinations are proposed herein.

***Pourthiaea blinii*** (Lévl.) Iketani & H.Ohashi subsp. ***podocarpifolia*** (T.T.Yu) B.B.Liu, **comb. nov.**

≡ *Photinia podocarpifolia* T.T.Yu, Acta Phytotax. Sin. 8: 230. 1963. ≡ *Pourthiaea podocarpifolia* (T.T.Yu) Iketani & H.Ohashi, J. Jap. Bot. 66(6): 354. 1991. Type: China. Guizhou: Luodian County, Bamao, Dating Xiang, Hongshui River, riverside, 200m, 8 April 1959, *Qiannan Expedition 408* (lectotype, designated by Liu & Hong (2016: 225): PE [barcode 00004600]!; isolectotypes HGAS [barcode 021090]!, KUN [barcode 606957, 747408]!, PE [barcode 00020608, 01498393, 01498394]!).

***Pourthiaea brevifolia*** (Cardot) B.B.Liu, **comb. nov.**

≡ *Photinia beauverdiana* C.K.Schneid. var. *brevifolia* Cardot, Notul. Syst. (Paris) 3: 378. 1918. ≡ *Pourthiaea beauverdiana* (C.K.Schneid.) Hatus. var. *Brevifolia* (Cardot) Iketani & H.Ohashi, J. Jap. Bot. 66(6): 353. 1991. Type: China. Western Hupeh [Hubei]: C. China, Shing-shan [Xingshan County] or Nanto, May 1900, *E.H. Wilson 794* (lectotype, designated by Liu & Hong (2017: 19): P [barcode P02143175]!; isolectotype: A [barcode 00045593]!).

***Pourthiaea dabeishanensis*** (M.B.Deng & G.Yao) B.B.Liu, **comb. nov.**

≡ *Photinia dabeishanensis* M.B.Deng & G.Yao, Bull. Nanjing Bot. Gard. 1984–1985: 126. 1986. Type: China. Anhui: Jinzhai County, Baimazhai Forest Farm, 1,100 m, 17 May 1984, *M.B. Deng 81741* (lectotype, designated by Liu & Hong (2017: 19): NAS [barcode NAS00072797]!; isolectotype: NAS [barcode NAS00072798]!).

***Pourthiaea magnoliifolia*** (Z.H.Chen) B.B.Liu, **comb. nov.**

≡ *Photinia magnoliifolia* Z.H.Chen, J. Zhejiang Forest. Coll. 3: 35. 1986. Type: CHINA. Zhejiang: Lin’an County, Qingshan Reservoir, 35 m, 15 April 1984, *Z.H. Chen & L.H. Ren 84006* HHBG!.

***Pourthiaea pustulata*** (Lindl.) B.B.Liu, **comb. nov.**

≡ *Photinia pustulata* Lindl., Edwards’s Bot. Reg. 23: sub t. 1956. 1837. ≡ *Pourthiaea arguta* subsp. *pustulata* (Lindl.) B.B.Liu & D.Y.Hong, Phytotaxa 325(1): 21. 2017. Type: China. “China prope Cantonem, Parkes” [Guangdong]. *s.coll. s.n.* (holotype: K [barcode K000758265]!).

II. ***Pourthiaea*** sect. ***Amphidoxae*** B.B.Liu, **sect. nov.** Type: *Pourthiaea amphidoxa* (C.K.Schneid.) Stapf

Diagnosis: This section could be easily distinguished from the other two sections in *Pourthiaea* with the following characters, two or five carpels, rarely four, never three, and larger fruits, usually more than 2 cm (*Pourthiaea amphidoxa* and *P. tomentosa*).

Distribution: Central and South China

Including three species, *Pourthiaea amphidoxa*, *P. pilosicalyx*, and *P. tomentosa*.

III. ***Pourthiaea*** sect. ***Impressivenae*** B.B.Liu, **sect. nov.** Type: *Pourthiaea impressivena* (Hayata) Iketani & H.Ohashi

Diagnosis: Characterized by ferruginous villous leaf blade, petiole, inflorescences, hypanthium, and branchlet, this section is distinct from others.

Distribution: East & South China.

Including three species, *Pourthiaea hirsuta*, *P. impressivena*, and *P. zhejiangensis*.

## CONCLUSIONS

In this study, we employed a genome skimming approach to reconstruct the phylogenetic relationships of *Pourthiaea*, utilizing both chloroplast genomes and a nrDNA dataset. Our analyses revealed significant topological incongruence between plastid and nuclear trees, spanning multiple species, and it challenged the monophyly of several widely distributed species. Notably, the nrDNA phylogeny aligned more closely with the species boundaries compared to the plastome phylogeny. Drawing upon evidence from morphological and phylogenomic analyses, we propose a new infrageneric classification for *Pourthiaea*, consisting of three sections.

Given the taxonomic challenges that no doubt await future researchers, it is imperative to make concrete plans to design research strategies that combine morphological, molecular, and ecological approaches to elucidate the taxonomy and phylogenetic relationships within the genus *Pourthiaea*. The recently proposed Deep Genome Skimming (DGS) method by Liu & al. (2021) offers the potential to assemble a vast number of single-copy nuclear genes for phylogenomic analyses. By integrating these diverse lines of evidence, taxonomists can work towards providing a more comprehensive and reliable classification system for this fascinating group of Rosaceae shrubs. These collective efforts will not only deepen our understanding of *Pourthiaea*’s evolutionary history, but also facilitate the development of more effective conservation strategies aimed at safeguarding these unique and ecologically important plant species.

## ACKNOWLEDGEMENTS

We express our gratitude to Wen-Bin Ju (Chengdu Institute of Biology, Chinese Academy of Sciences), En-De Liu (Kunming Institute of Botany, Chinese Academy of Sciences), Guan-Ling Sun (Conghua, Guangdong, China), Long-Yuan Wang (Zhongkai University of Agriculture and Engineering), Lei Xie (Beijing Forestry University), and Xin-Tang Ma and Qin Ban (Institute of Botany, Chinese Academy of Sciences) for their valuable contributions in sample collections. We would also like to extend our thanks to Ze-Long Nie (Jishou University), Chen Ren (South China Botanical Garden, Chinese Academy of Sciences), Wen-Pan Dong (Beijing Forestry University) for their assistance with data analyses, and Jing Xuan (Institute of Botany, Chinese Academy of Sciences) for his efforts in preparing the sampling map. The computations carried out in this study were performed on the Dell T7920 workstation, owned by Bin-Bin Liu (Institute of Botany, Chinese Academy of Sciences), and the Smithsonian High Performance Cluster (SI/HPC) at the Smithsonian Institution (https://doi.org/10.25572/SIHPC). Financial support for this work was provided by the National Natural Science Foundation of China, grant number 32270216 (to BBL) and 32000163 (to BBL), as well as the Youth Innovation Promotion Association CAS, grant number 2023086 (to BBL).

## AU THOR CONTRIBUTIONS

BBL designed and supervised this study. BBL and CX conducted the experiment, and GNL, DKM, YZ, SYX, HW, ZTJ, and FXL analyzed the data. GNL wrote the early version of the manuscript; RGJH, SHJ and LZ provided suggestions for structuring the manuscript; BBL and YW provided comments on structuring the manuscript and contributed to the discussion and revision of the manuscript. All authors approved the final manuscript. — GNL, https://orcid.org/0009-0009-0765-0392; DKM, https://orcid.org/0009-0005-5523-508X; YZ, https://orcid.org/0000-0002-0802-3923; RGJH, https://orcid.org/0000-0002-2896-4907; SYX, https://orcid.org/0000-0002-0227-0982; HW, https://orcid.org/0009-0009-9075-698X; ZTJ, https://orcid.org/0000-0003-1358-0043; FXL, https://orcid.org/0000-0003-1097-9438; SHJ, https://orcid.org/0000-0003-0334-6683; LZ, https://orcid.org/0000-0003-3846-505X; CX, https://orcid.org/0000-0002-9678-4772; YW, https://orcid.org/0000-0002-2013-4829; BBL, https://orcid.org/0000-0002-0297-7531

## SUPPORTING INFORMATION

Supporting Information may be found online in the Supporting Information section at the end of the article.

**Fig. S1** Gene map of the *Pourthiaea* chloroplast genome. The genes inside and outside of the circle are transcribed in the clockwise and counterclockwise directions, respectively. Genes belonging to different functional groups are shown in different colors. The thick lines indicate the extent of the inverted repeats (IRa and IRb) that separate the genomes into small single-copy (SSC) and large single-copy (LSC) regions.

**Fig. S2** Maximum likelihood phylogeny of *Pourthiaea* in the framework of Maleae inferred from RAxML analysis of the nuclear ribosomal DNA (nrDNA) data matrix. Numbers above the branches indicate Bootstrap Support (BS).

**Fig. S3** Maximum likelihood phylogeny of *Pourthiaea* in the framework of Maleae inferred from IQ-TREE2 analysis of the nuclear ribosomal DNA (nrDNA) data matrix. Numbers above the branches indicate the SH-aLRT support and Ultrafast Bootstrap support.

**Fig. S4** Bayesian inference tree of *Pourthiaea* in the framework of Maleae based on the nuclear ribosomal DNA (nrDNA) data matrix by MrBayes 3.2.7a. Numbers above the branches indicate the BI posterior probabilities.

**Fig. S5** Maximum likelihood phylogeny of *Pourthiaea* with one outgroup (*Cydonia oblonga*) inferred from RAxML analysis of the whole plastome data matrix. Numbers above the branches indicate Bootstrap Support (BS).

**Fig. S6** Maximum likelihood phylogeny of *Pourthiaea* with one outgroup (*Cydonia oblonga*) inferred from IQ-TREE2 analysis of the whole plastome data matrix. Numbers above the branches indicate the SH-aLRT support and Ultrafast Bootstrap support.

**Fig. S7** Bayesian inference tree of *Pourthiaea* with one outgroup (*Cydonia oblonga*) based on the whole plastome data matrix by MrBayes 3.2.7a. Numbers above the branches indicate the BI posterior probabilities.

**Fig. S8** Maximum likelihood phylogeny of *Pourthiaea* with one outgroup (*Cydonia oblonga*) inferred from RAxML analysis of the 79 concatenated plastid coding genes data matrix. Numbers above the branches indicate Bootstrap Support (BS).

**Fig. S9** Maximum likelihood phylogeny of *Pourthiaea* with one outgroup (*Cydonia oblonga*) inferred from IQ-TREE2 analysis of the 79 concatenated plastid coding genes data matrix. Numbers above the branches indicate the SH-aLRT support and Ultrafast Bootstrap support.

**Fig. S10** Bayesian inference tree of *Pourthiaea* with one outgroup (*Cydonia oblonga*) based on the 79 concatenated plastid coding genes data matrix by MrBayes 3.2.7a. Numbers above the branches indicate the BI posterior probabilities.

**Fig. S11** Species tree of *Pourthiaea* with one outgroup (*Cydonia oblonga*) inferred from ASTRAL-III of the concatenated 79 plastid coding genes. Numbers above the branches indicate the branch support values measuring the support for a local posterior possibility.

**Fig. S12** Maximum likelihood phylogeny of *Pourthiaea* in the framework of Maleae inferred from RAxML analysis of the whole plastome data matrix. Numbers above the branches indicate Bootstrap Support (BS).

**Fig. S13** Maximum likelihood phylogeny of *Pourthiaea* in the framework of Maleae inferred from IQ-TREE2 analysis of the whole plastome data matrix. Numbers above the branches indicate the SH-aLRT support and Ultrafast Bootstrap support.

**Fig. S14** Bayesian inference tree of *Pourthiaea* in the framework of Maleae based on the whole plastome data matrix by MrBayes 3.2.7a. Numbers above the branches indicate the BI posterior probabilities.

**Fig. S15** Maximum likelihood phylogeny of *Pourthiaea* in the framework of Maleae inferred from RAxML analysis of the 79 concatenated plastid coding genes data matrix. Numbers above the branches indicate Bootstrap Support (BS).

**Fig. S16** Maximum likelihood phylogeny of *Pourthiaea* in the framework of Maleae inferred from IQ-TREE2 analysis of the 79 concatenated plastid coding genes data matrix. Numbers above the branches indicate the SH-aLRT support and Ultrafast Bootstrap support.

**Fig. S17** Bayesian inference tree of *Pourthiaea* in the framework of Maleae based on the 79 concatenated plastid coding genes data matrix by MrBayes 3.2.7a. Numbers above the branches indicate the BI posterior probabilities.

**Fig. S18** Species tree of *Pourthiaea* in the framework of Maleae inferred from ASTRAL-III of the concatenated 79 plastid coding genes. Numbers above the branches indicate the branch support values measuring the support for a local posterior possibility.

