## Appendix 1 for "Phylogenomic analyses support a new infrageneric classification of *Pourthiaea* (Maleae, Rosaceae) using multiple inference methods and extensive taxon sampling"

**Appendix 1.** Accessions of *Pourthiaea* used in this study.

The list of GenBank accessions used in this study is presented as follows: Taxon, unique identifier, collector and collection number for the voucher (Herbarium), collection locality, BioProject, SRA accession, GenBank accession number for chloroplast genome, chloroplast length (bp), GenBank accession number for nrDNA, nrDNA length (bp). A dash (–) indicates data that were unavailable. Sequences generated for this study are marked with an asterisk (^*^). The designations of α, β, and γ were indicated in the materials and methods.

***Amelanchier sinica*** (Schneid.) Chun, –, *B.B.Liu & G.N.Liu 220725* (PE), China: Beijing, PRJNA952323, SRR24154176, MK920291^*^, 159901, MN216016^*^, 6409; ***Crataegus marshallii*** Eggl., –, *J.B.Nelson#26961* (US-03700739), USA, PRJNA952323, SRR24154175, MK920293^*^, 159660, MN215977^*^, 6459; ***Cydonia oblonga*** Mill., –, *B.B.Liu & G.N.Liu#3873* (PE), China: Beijing, PRJNA952323, SRR24154163, MN061993^*^, 159643, MN216014^*^, 6234; ***Gillenia trifoliata*** (L.) Moench, –, *B.B.Liu#4677* (US), USA: Washionton DC, PRJNA952323, SRR24154152, MN068252^*^, 159400, MN577923^*^, 6351; ***Hesperomeles glabrata*** (Kunth) M.Roem., –, *P.E.Berry#4561* (US-03695884), Venezuela, PRJNA952323, SRR24154141, MK920298^*^, 160176, MN216003^*^, 6469; ***Kageneckia oblonga*** Ruiz & Pav., –, *M.Mahu & C.H.Badilla#*10358 (US-03694604), Chile: Maule Chanco, PRJNA952323, SRR24154130, MN068266^*^, 159409, MN577932^*^, 6340; ***Lindleya mespiloides*** Kunth, –, *G.B.Hinton & al.#18696* (US-00903453), Mexico, PRJNA952323, SRR24154119, MN068248^*^, 158697, MN577906^*^, 6337; ***Malus baccata*** (L.) Borkh., –, *J.Wen#*14050 (US), USA: Minnesota, PRJNA952323, SRR24154109, MK896774^*^, 160024, MN215980^*^, 6413; ***Phippsiomeles matudae*** B.B.Liu & J.Wen, –, *J.B.Phipps & P.G.Smith#5865* (US-00909013), Mexico, PRJNA952323, SRR24154108, MN062002^*^, 159668, MN216001^*^, 6420; ***Photinia glabra*** (Thunb.) Maxim., –, *B.B.Liu#P1901-2* (PE-02071209), China: Guangxi, PRJNA952323, SRR24154172, MK920277^*^, 159571, MN216020^*^, 6603; ***Pourthiaea amphidoxa*** (C.K.Schneid.) Stapf, P1, *B.B.Liu#2072* (PE)^α^, China: Hubei, PRJNA952323, SRR24154174, MT249042^*^, 160244, MT249083^*^, 6328; ***Pourthiaea amphidoxa*** (C.K.Schneid.) Stapf, P2, *W.B.Ju & H.N.Deng#HGX12711* (CDBI-CDBI0227127)^α^, China: Sichuan, PRJNA952323, SRR24154173, MT249049^*^, 160344, MT249090^*^, 6328; ***Pourthiaea amphidoxa*** (C.K.Schneid.) Stapf, P3, *Y.F.Deng* *& al.#11700* (PE-01598124)^α^, China: Hunan, PRJNA952323, SRR24154171, MT249052^*^, 160149, MT249093^*^, 6330; ***Pourthiaea amphidoxa*** (C.K.Schneid.) Stapf, P4, *S.Q.Chen#16816* (PE-00738651)^α^, China: Guangxi, PRJNA952323, SRR24154170, MT249053^*^, 160285, MT249094^*^, 6328; ***Pourthiaea amphidoxa*** (C.K.Schneid.) Stapf, P5, *B.B.Liu#P1918-4* (PE-02090670)^β^, China: Chongqing, PRJNA952323, SRR24154169, MT249055^*^, 160279, MT249096^*^, 6328; ***Pourthiaea amphidoxa*** (C.K.Schneid.) Stapf, P6, *B.B.Liu#2530* (PE)^β^, China: Guizhou, PRJNA952323, SRR24154168, MT249059^*^, 160397, MT249100^*^, 6329; ***Pourthiaea amphidoxa*** (C.K.Schneid.) Stapf, P7, *L.Xie#4620* (PE-02050150)^β^, China: Sichuan, PRJNA952323, SRR24154167, MN061992^*^, 160357, MN216012^*^, 6328; ***Pourthiaea amphidoxa*** (C.K.Schneid.) Stapf, P8, *M.T.An#0086* (PE-00738553)^β^, China: Guizhou, PRJNA952323, SRR24154166, MT249061^*^, 160195, MT249102^*^, 6328; ***Pourthiaea amphidoxa*** (C.K.Schneid.) Stapf, P9, *Huaping Nature Reserve Exp.#H0528* (PE-02108460)^β^, China: Guangxi, PRJNA952323, SRR24154165, MT249062^*^, 160341, MT249103^*^, 6329; ***Pourthiaea arguta*** (Lindl.) Decne., P22, *J.Wen & al.#11063* (US-00863110)^γ^, Vietnam: Lam Dong, PRJNA952323, SRR24154164, OR149024^*^,160634, OR162028^*^, 6335; ***Pourthiaea beauverdiana*** (C.K.Schneid.) Hatus., P10, *B.B.Liu#2446* (PE-02080232)^α^, China: Jiangxi, PRJNA952323, SRR24154162, MT249027^*^, 160288, MT249068^*^, 6328; ***Pourthiaea beauverdiana*** (C.K.Schneid.) Hatus., P11, *B.B.Liu#2124* (PE-02080624)^α^, China: Guangdong, PRJNA952323, SRR24154161, MT249028^*^, 160342, MT249069^*^, 6328; ***Pourthiaea beauverdiana*** (C.K.Schneid.) Hatus., P12, *B.B.Liu#2223* (PE-02080202)^α^, China: Hubei, PRJNA952323, SRR24154160, MT249035^*^, 160309, MT249076^*^, 6328; ***Pourthiaea beauverdiana*** (C.K.Schneid.) Hatus., P13, *B.B.Liu#P1911-1* (PE-02080334)^α^, China: Guizhou, PRJNA952323, SRR24154159, MT249037^*^, 160280, MT249078^*^, 6329; ***Pourthiaea beauverdiana*** (C.K.Schneid.) Hatus., P17, *B.B.Liu#2448* (PE-02068099)^α^, China: Hunan, PRJNA952323, SRR24154158, MT249046^*^, 160343, MT249087^*^, 6328; ***Pourthiaea beauverdiana*** (C.K.Schneid.) Hatus., P18, *E.D.Liu#LED6340* (KUN)^α^, China: Yunnan, PRJNA759205, SRR15691188, MT249048, 160284, MT249089, 6328; ***Pourthiaea beauverdiana*** (C.K.Schneid.) Hatus., P20, *B.B.Liu#2208* (PE-02080332)^β^, China: Sichuan, PRJNA952323, SRR24154157, MN061991^*^, 160159, MN216011^*^, 6328; ***Pourthiaea beauverdiana*** (C.K.Schneid.) Hatus., P21, *Z.B.Jian & al.#50711* (PE-00337378)^β^, China: Guizhou, PRJNA952323, SRR24154156, MT249067^*^, 160337, MT249108^*^, 6328; ***Pourthiaea blinii*** subsp. ***blinii*** (H.Lév.) Iketani & H.Ohashi, P25, *B.B.Liu#2171* (PE-02070183)^α^, China: Guizhou, PRJNA952323, SRR24154155, MT249034^*^, 160391, MT249075^*^, 6328; ***Pourthiaea blinii*** subsp. ***blinii*** (H.Lév.) Iketani & H.Ohashi, P27, *S.S.Zhou#1907* (PE-01498581)^β^, China: Yunnan, PRJNA952323, SRR24154154, MT249060^*^, 160332, MT249101^*^, 6329; ***Pourthiaea blinii*** subsp. ***blinii*** (H.Lév.) Iketani & H.Ohashi, P28, *S.D.Zhang#sd023*, China: Yunnan, SRR24154151, KY419919^*^, 133404, –, –; ***Pourthiaea blinii*** subsp. ***podocarpifolia*** (T.T.Yu) Iketani & H.Ohashi, P26, *B.B.Liu#2158* (PE-02070120)^β^, China: Guangxi, PRJNA952323, SRR24154153, MN061990^*^, 160307, MN216010^*^, 6328; ***Pourthiaea brevifolia***, P15, *B.B.Liu#2227* (PE-02080198)^α^, China: Hubei, PRJNA952323, SRR24154150, MT249039^*^, 160312, MT249080^*^, 6329; ***Pourthiaea dabeishanensis***, P16, *B.B.Liu#2325* (PE-02080098)^α^, China: Anhui, PRJNA952323, SRR24154149, MT249044^*^, 160269, MT249085^*^, 6328; ***Pourthiaea hirsuta*** (Hand.-Mazz.) Iketani & H.Ohashi, P29, *B.B.Liu#2385* (PE)^α^, China: Zhejiang, PRJNA952323, SRR24154147, MT249031^*^, 160255, MT249072^*^, 6328; ***Pourthiaea hirsuta*** (Hand.-Mazz.) Iketani & H.Ohashi, P30, *B.B.Liu#2595* (PE)^α^, China: Fujian, PRJNA952323, SRR24154148, MN061986^*^, 160233, MN215983^*^, 6328; ***Pourthiaea impressivena*** (Hayata) Iketani & H.Ohashi, P31, *G.L.Sun#180* (PE)^α^, China: Guangdong, PRJNA952323, SRR24154146, MT249047^*^, 160200, MT249088^*^, 6328; ***Pourthiaea lucida*** Decne., P19, *C.M.Wang#53078* (PE-01515399)^α^, China: Taiwan, PRJNA952323, SRR24154145, MT249051^*^, 160269, MT249092^*^, 6329; ***Pourthiaea magnoliifolia*** (Lindl.) Decne., P14, *B.B.Liu#P1953-1* (PE-02070470)^α^, China: Zhejiang, PRJNA952323, SRR24154144, MT249038^*^, 160318, MT249079^*^, 6329; ***Pourthiaea parvifolia*** E.Pritz., P39, *B.B.Liu#P2083-1* (PE-02071678)^α^, China: Hubei, PRJNA952323, SRR24154143, MT249029^*^, 160283, MT249070^*^, 6329; ***Pourthiaea parvifolia*** E.Pritz., P40, *B.B.Liu#2587* (PE-02099781)^α^, China: Fujian, PRJNA952323, SRR24154142, MT249030^*^, 160367, MT249071^*^, 6331; ***Pourthiaea parvifolia*** E.Pritz., P41, *B.B.Liu#2418* (PE-02099829)^α^, China: Fujian, PRJNA952323, SRR24154140, MT249032^*^, 160293, MT249073^*^, 6329; ***Pourthiaea parvifolia*** E.Pritz., P42, *B.B.Liu#P1914-3* (PE-02071426)^α^, China: Guizhou, PRJNA952323, SRR24154139, MT249036^*^, 160283, MT249077^*^, 6329; ***Pourthiaea parvifolia*** E.Pritz., P44, *B.B.Liu#P1906-1* (PE-02070280)^α^, China: Guangxi, PRJNA952323, SRR24154138, MT249041^*^, 160142, MT249082^*^, 6328; ***Pourthiaea parvifolia*** E.Pritz., P45, *B.B.Liu#P1975-1* (PE-02070711)^α^, China: Jiangxi, PRJNA952323, SRR24154137, MT249043^*^, 160390, MT249084^*^, 6328; ***Pourthiaea parvifolia*** E.Pritz., P47, *B.B.Liu#P1919-3* (PE-02071296)^β^, China: Chongqing, PRJNA952323, SRR24154136, MN061989^*^, 160401, MN216009^*^, 6328; ***Pourthiaea parvifolia*** E.Pritz., P50, *B.B.Liu#P1967-4* (PE-02069810)^β^, China: Zhejiang, PRJNA952323, SRR24154135, MT249058^*^, 160496, MT249099^*^, 6328; ***Pourthiaea parvifolia*** E.Pritz., P53, *C.C.Liao & al.#530* (US-03699539)^γ^, China: Taiwan, PRJNA952323, SRR24154134, OR149023, 160271, OR162027, 6335; ***Pourthiaea pilosicalyx*** (T.T.Yu) Iketani & H.Ohashi, P32, *B.B.Liu#2131* (PE-02080439)^α^, China: Guangxi, PRJNA759205, SRR15691186, MT249045, 160142, MT249086, 6330; ***Pourthiaea pilosicalyx*** (T.T.Yu) Iketani & H.Ohashi, P33, *G.F.Wang#29437* (PE-01498390)^α^, China: Guizhou, PRJNA952323, SRR24154133, MN216024^*^, 160330, MN216008^*^, 6328; ***Pourthiaea pustulata***, P23, *B.B.Liu#2105* (PE-02080663)^α^, China: Hainan, PRJNA952323, SRR24154132, MT249033^*^, 160258, MT249074^*^, 6329; ***Pourthiaea pustulata***, P24, *B.B.Liu#213502* (PE)^α^, China: Guangdong, PRJNA952323, SRR24154131, MT249054^*^, 160310, MT249095^*^, 6330; ***Pourthiaea sorbifolia*** (W.B.Liao & W.Guo) B.B.Liu & D.Y.Hong, P35, *Wuling Expedition#909* (PE-01364106)^β^, China: Hunan, PRJNA952323, SRR24154129, MN061994^*^, 160371, MN216017^*^, 6328; ***Pourthiaea sorbifolia*** (W.B.Liao & W.Guo) B.B.Liu & D.Y.Hong, P36, *Wulingshan Expedition Team#2302* (PE-01896033)^β^, China: Hunan, PRJNA952323, SRR24154128, MT249065^*^, 160207, MT249106^*^, 6330; ***Pourthiaea tomentosa*** (T.T.Yu & T.C.Ku) B.B.Liu & J.Wen, P37, *G.F.Li#60885* (PE-00739363)^β^, China: Chongqing, PRJNA952323, SRR24154127, MT249066^*^, 160337, MT249107^*^, 6328; ***Pourthiaea tomentosa*** (T.T.Yu & T.C.Ku) B.B.Liu & J.Wen, P38, *B.B.Liu#4674* (PE)^β^, China: Chongqing, PRJNA952323, SRR24154126, MN061995^*^, 160290, MN216018^*^, 6328; ***Pourthiaea tsaii*** (Rehder) H. Iketani et Ohashi, P34, *B.B.Liu#2186* (PE-02080401)^α^, China: Yunnan, PRJNA952323, SRR24154125, MN061987^*^, 160199, MN215993^*^, 6328; ***Pourthiaea villosa*** (Thunb.) Decne., P43, *B.B.Liu#P1978-1* (PE-02070057)^α^, China: Jiangxi, PRJNA952323, SRR24154124, MT249040^*^, 160231, MT249081^*^, 6329; ***Pourthiaea villosa*** (Thunb.) Decne., P46, *K.Yonekura#1907* (PE-01523677)^α^, Japan: Honshu, PRJNA952323, SRR24154123, MT249050^*^, 1603600, MT249091^*^, 6329; ***Pourthiaea villosa*** (Thunb.) Decne., P48, *B.B.Liu#P2085-2* (PE-02071412)^β^, China: Hubei, PRJNA952323, SRR24154122, MT249056^*^, 160432, MT249097^*^, 6329; ***Pourthiaea villosa*** (Thunb.) Decne., P49, *B.B.Liu#P1993-1* (PE-02070879)^β^, China: Shandong, PRJNA952323, SRR24154121, MT249057^*^, 160352, MT249098^*^, 6329; ***Pourthiaea villosa*** (Thunb.) Decne., P51, *M.Furuse#50324* (PE-01656471)^β^, Japan: Honshu, PRJNA952323, SRR24154120, MT249063^*^, 160446, MT249104^*^, 6329; ***Pourthiaea villosa*** (Thunb.) Decne., P52, *M.Furuse#29437* (PE-01156868)^β^, Japan: Hokkaido, PRJNA952323, SRR24154118, MT249064^*^, 160259, MT249105^*^, 6330; ***Pourthiaea zhejiangensis*** (P.L.Chiu) Iketani & H.Ohashi, P54, *L.Y.Wang#T-29509* (SYS)^α^, China: Zhejiang, PRJNA759205, SRR15691185, MN061988, 160300, MN216007, 6328; ***Rhaphiolepis cavaleriei*** (H.Lév.) B.B.Liu & J.Wen, –, *B.B.Liu#2585* (PE-02070509), China: Hunan, PRJNA952323, SRR24154117, MK920283^*^, 159210, MN215982^*^, 6450; ***Sorbus commixta*** Hedl., –, *M.Li#HG019* (CDBI), South Korea, PRJNA952323, SRR24154116, MK920288^*^, 159952, MN215997^*^, 6804; ***Sorbus tianschanica*** Rupr., –, *M.Li#zzm848* (CDBI), China, PRJNA952323, SRR24154115, MK920289^*^, 160037, MN215998^*^, 6410; ***Sorbus ulleungensis*** Chin S.Chang, –, –, South Korea, –, MG011706, 159632, –, –; ***Sorbus vilmorinii*** Schneid., –, *Y.S.Chen & al.#13-1059* (PE), China, PRJNA952323, SRR24154114, MK920285^*^, 159936, MN215994^*^, 6409; ***Vauquelinia australis*** Standl., *W.Hess & G.Wilhelm#4382* (US-00908940), Mexico, PRJNA952323, SRR24154113, MN068250^*^, 159813, MN577917^*^, 6337; ***Vauquelinia californica*** subsp. ***sonorensis*** W.J.Hess & Henr., *J.Henrickson#20281* (US-00903444), Mexico, PRJNA952323, SRR24154112, MN068269^*^, 158894, MN905593^*^, 5778; ***Vauquelinia corymbosa*** subsp. ***heterodon*** (I.M.Johnst.) W.J.Hess & Henr., *J.Henrickson#189230* (US-00903439), Mexico, PRJNA952323, SRR24154111, MN068249^*^, 159857, MN905576^*^, 5778; ***Vauquelinia pauciflora*** Standl., *F.Reichenbacher#872* (US-03694614), USA, PRJNA952323, SRR24154110, MN068251^*^, 159772, MN905577^*^, 5858.
